# Host evolution improves genetic circuit function in complex growth environments

**DOI:** 10.1101/2024.03.13.583595

**Authors:** Joanna T. Zhang, Andrew Lezia, Philip Emmanuele, Muyao Wu, Connor A. Olson, Adam M. Feist, Jeff Hasty

## Abstract

Genetically engineered bacteria have become an attractive platform for numerous biomedical and industrial applications. Despite genetic circuitry functioning predictably under favorable growth conditions in the lab, the same cannot be said when placed in more complex environments for eventual deployment. Here, we used a combination of evolutionary and rational engineering approaches to enhance *E. coli* for robust genetic circuit behavior in non-traditional growth environments. We utilized adaptive laboratory evolution (ALE) on *E. coli* MG1655 in a minimal media with a sole carbon source and saw improved dynamics of a population-lysis-based circuit after host evolution. Additionally, we improved lysis circuit tolerance of a more clinically relevant strain, the probiotic *E. coli* Nissle, using ALE of the host strain in a more complex media environment with added reactive oxygen species (ROS) stress. We observed improved recovery from circuit-induced lysis in the evolved Nissle strain, and in combination with directed mutagenesis, recovered circuit function in the complex media. These findings serve as a proof-of-concept that relevant strains of bacteria can be optimized for improved growth and performance in complex environments using ALE and that these changes can modify and improve synthetic gene circuit function for real-world applications.

## Introduction

As genetic engineering tools improve in sophistication, synthetic biologists are able to develop increasingly complex genetic circuits for control of microbial population behavior (*1–3*). These circuits have been used for the detection and treatment of cancer (*4, 5*), mitigation of metabolic disorders (*6*), and sensing of the gut environment (*7*). However, numerous limitations remain in the real-world deployment of engineered gene circuits, such as unpredictability of circuit behavior in complex growth environments (*8*). Quantitative modelling of physiological transitions in *E. coli* from one growth medium to another showed marked differences in metabolic activity in response to changes in medium nutrients, which can significantly influence protein production and affect underlying circuits (*9,10*). Expression of synthetic gene cassettes further places stress on engineered populations, increasing sensitivity to external disturbances (*11*). Thus, gene circuits that behave consistently in nutrient-rich growth conditions in the lab can oftentimes break down when transferred to more complex and non-traditional environments (*12*).

Traditional tools to improve genetic circuit behavior employ rational engineering principles, where individual circuit components are chosen to meet specific requirements. For example, mutant libraries can be generated and screened for functionality, a process that may be time-consuming and heavily relies on trial-and-error analysis and testing (*13, 14*). Control engineering principles can also be employed to limit the influence of external perturbations on circuit function (*15*). However, these often require multiple genetic components that place significant metabolic burden on the cells. Recently, the behavior of the toggle switch was quantitatively modelled in *E. coli* in natural, fluctuating environments, but translation to more complex circuits in other strains remain difficult (*16*).

Adaptive laboratory evolution (ALE), where a candidate strain is cultured for a prolonged period under a selective pressure, enables the accumulation of genetic mutations for improved growth in that environment over time (*17*). This technique has been used in both bacteria and yeast for the generation of desired phenotypes such as osmotic, acid, and temperature tolerance, and increased alternate substrate metabolism for bioproduction (*18*). Recently, ALE was used to optimize growth performance of a genome-reduced *E. coli* strain (*19*), co-evolve mutually auxotrophic strains of yeast and lactic acid bacteria for improved production of the auxotrophic compound (*20*), and improve methanol utilization in *E. coli* engineered with a hybrid methanol assimilation pathway (*21*). In combination with omics technologies, researchers can better understand mutational pathways underlying evolution and create strains that are tolerant to unique and stressful environmental conditions (*22*). Genetic circuitry has also been incorporated with adaptive evolution, where the mutation rate is directly controlled by the concentration of a target metabolite in a feedback loop, to increase tyrosine and isoprenoid production (*23*).

To date, host evolutionary engineering has been under-utilized for the improvement of dynamic genetic circuits in desired environments. In this study, we leveraged the selected genetic changes from evolved strains as an alternative method of improving circuit dynamics in complex environments. As a proof of concept, we first applied selective pressure on wild-type *E. coli* K-12 MG1655 in a simple media and found improved growth in the media during the evolution. We chose a previously developed Synchronized Lysis Circuit (SLC), which exhibits dynamic growth rate dependent behavior with potential *in vivo* applications, as our candidate circuit for evaluation (*3, 4, 24*). The SLC causes cycles of population lysis and regrowth and we observed improvements in circuit behavior and burden tolerance after ALE. Next, we directly targeted SLC tolerance of *E. coli* Nissle, a more clinically relevant strain, through evolution of the host strain to Paraquat (PQ)-induced reactive oxygen species (ROS) stress and saw improved growth under PQ stress and recovery from circuit induced lysis. We further increased the complexity of this evolution by using a physiologically relevant media base. Using directed mutagenesis on the evolved Nissle, in combination with batch culture and high throughput microfluidic screening, we identified a fully oscillatory strain in the complex media, thus showing the synergy of ALE with traditional synthetic biology tools. Finally, we explored the differing host-circuit effects between wildtype MG1655 and Nissle using a ribosome binding site (RBS) library and uncovered important differences between the two strains. Overall, this work highlights the utility of combining evolutionary and rational engineering to produce robust genetic circuits, as well as providing a framework for translation of engineered circuits from *E. coli* MG1655 to Nissle.

## Results

### Adaptive Laboratory Evolution In Simple Media Improved Genetic Circuit Behavior in MG1655

As an initial proof-of-concept, wild-type *E. coli* K-12 MG1655 was evolved in M9, a minimal media, with lactate as the sole carbon source. Six replicates were evolved in parallel (Fig 1a). In batch culture growth and passage, we observed that the growth rate on M9 lactate increased with subsequent evolutionary passages in all replicates (Fig 1b). An individual clone from each replicate evolution was sequenced for mutations at the end of the evolution (Table S1). We identified convergent mutations in *ppsA* (phosphoenolpyruvate synthase in the gluconeogenesis pathway) and *ilvH* (acetolactate synthase subunit, both which were found to be mutated in previous ALE experiments in M9 lactate (*25–27*). *rhsE*, a pseudogene, was also identified as a convergent mutation but was likely a result of a non-isogeneic starting strain stock (Table S1) (*28*). As the ALE16 lineage contained representative mutations and similar growth rates to other lineages, clones from this replicate at multiple stages of the evolution were chosen for further analysis of circuit behavior.

**Figure 1:**
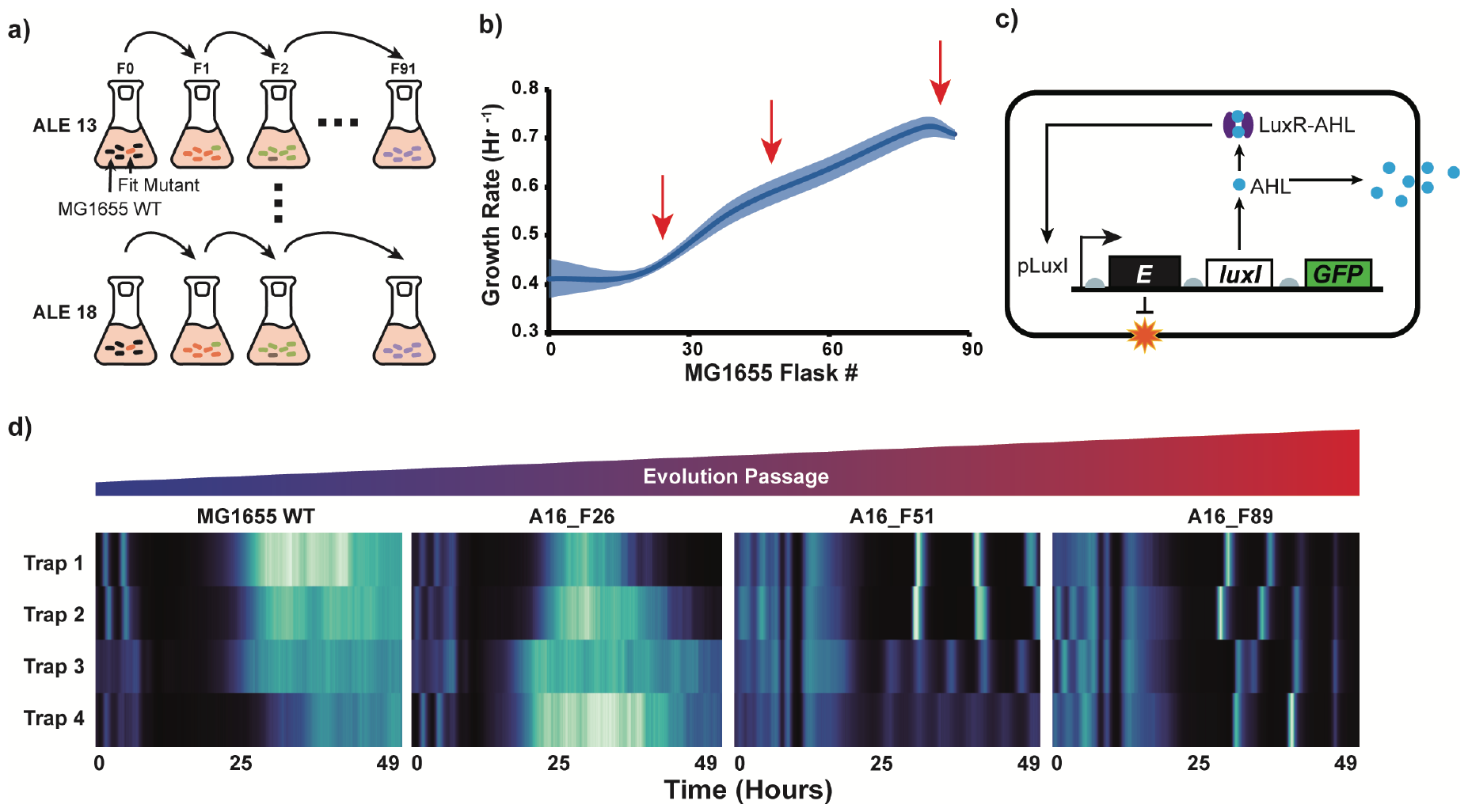
Evolution and characterization of MG1655 strains on M9 lactate. **a)** Schematic of ALE process. MG1655 wildtype strains were evolved in M9 lactate in 6 parallel replicate experiments (n = 6). F indicates flask or batch number **b)** Average growth rate of the similarly performing replicates over the course of evolution. Red arrows indicate flasks that were chosen for further analysis. All data points represent mean (solid line) ± SEM (shaded areas), n = 6. **c)** Diagram of the single plasmid synchronized lysis circuit (SLC). AHL production from LuxI synchronizes cells within the population and provides positive feedback. The X174E protein causes cell lysis as negative feedback. **d)** Heatmaps representing GFP time traces (proxy for circuit activation) of the MG1655 wildtype and different evolved strains from one lineage with the SLC.

To study the effect of the media-optimized evolved host strain on circuit response, we chose to characterize the synchronized lysis circuit (SLC), as it requires the combination of multiple circuit components and is sensitive to environmental factors and growth rate. In the SLC, expression of the LuxI protein leads to production of quorum sensing autoinducer N-Acyl homoserine lactone (AHL) for synchronous population behavior. Upon reaching a threshold population size and AHL concentration, the pLux promoter drives expression of the lysis protein E from phage X174 (X174E), leading to synchronized population lysis. Following lysis, a few cells survive and reseed the population, leading to continuous cycles of growth and lysis. We transformed a single plasmid version of the SLC into wildtype MG1655, ALE16-Flask26 (A16-F26), A16-F51, and A16-F89 (Fig 1c). In order to eliminate inter-experimental variations, we chose to use a multi-strain microfluidic device that can accommodate up to 48 unique strains of *E. coli* for testing (Fig S1). Strains in each trap were grown up in LB overnight, followed by a media change to M9 lactate. During the media change, all strains continued to oscillate. As M9 lactate completely replaced LB, both wildtype MG1655 and A16-F26 exhibited one last lysis peak at 7 hours followed by a prolonged delay before regrowth without continued lysis. Meanwhile, both A16-51 and A16-89 regained consistent lysis dynamics following the media change (Fig 1d, Video S1). Further analysis showed F51 and F89 shared mutations in *ppsA* as well as *hfq*, which plays a role in stress response (Table S1).

We hypothesized that the improvement in circuit dynamics at A16-F51 and A16-F89 was due to increased tolerance of the host strain in M9 lactate from the evolution. As production of the X174E protein at quorum caused population lysis, the wildtype and F26 strains were unable to recover due to suboptimal metabolism and growth in M9 lactate. Meanwhile, the accumulated genetic mutations at F51 and F89 allowed for enhanced metabolic activity in the M9 lactate environment. This led to better recovery from lysis, allowing continuous cycles of growth and lysis as seen in microfluidics (Fig 1d, Video S1). Further, when we compared promoter sensitivity in M9 lactate batch culture by testing each strain with a plasmid with *pLux* promoter driving GFP expression, we saw that A16-F89 showed inducibility to 1nM AHL whereas wildtype did not respond (Fig S2a). Interestingly, we also noticed that upon AHL induction, the wildtype strain exhibited a greater reduction in growth compared to A16-F89 (Fig S2b). Since GFP expression also places burden on the cells, it appears that the evolved strain could better handle this burden in M9 lactate.

### Evolution in Paraquat Improved *E. coli* Nissle Tolerance to Lysis Circuit Burden

Although *E. coli* K-12 MG1655 is widely used for development and testing of new synthetic circuits due to its favorable growth rate and easy transformability, it becomes a less suitable host candidate when considering therapeutic applications. Among various bacterial strains explored, *E. coli* Nissle, a probiotic currently in use for treatment of gastrointestinal disorders, has shown promise in clinical applications due to its favorable safety profile and relevance in areas such as irritable bowel disease (IBD), toxic gut metabolite accumulation, and in particular, treatment and detection of cancers (*29–32*). As such, we sought to use ALE to improve circuit behavior in *E. coli* Nissle in a more complex and physiologically relevant growth environment. We chose DMEM, a media formulation used for culturing mammalian cells, for our base media. The media was additionally modified to include a pH of 6.5, 0.4mM glucose, and 30mM lactate in order to roughly mimic stressors found in the tumor microenvironment (*33*). As we noticed that ALE improved circuit tolerance in the proof-of-concept study in MG1655 (previous section), we hypothesized that Nissle could be directly adapted for faster recovery of survivors following SLC lysis. Recent work shows that the X174E protein prevents peptidoglycan synthesis (*34*), which can lead to production of reactive oxygen species (ROS) stress (*35*). Thus, we chose to also include 350uM of Paraquat (PQ), a compound that generates ROS.

Five Nissle ALE replicates were evolved in parallel to generate an optimized host strain for evaluation of the SLC. We observed an increase in the evolved Nissle lineage’s growth rate within the first 30 passages, after which the rate began to plateau (Fig S3a). Each replicate lineage was sequenced for mutations at the end of the evolution in order to capture the adaptive mutational response. We found convergent mutations in *argP*, which controls transcription of genes related to arginine and lysine metabolism, suggesting evolution to the growth media base, and *rpoA*, a transcriptional subunit that was found to affect anaerobically expressed genes in *Salmonella* (*36*) (Table S1). These genes are different from a previous study utilizing ALE and multiomics analysis for the impact of PQ stress on *E. coli* MG1655 physiology, most likely due to a lower PQ concentration, a significantly different evolution growth medium (i.e., a complex medium in this case), and a different strain subtype (*37*). As the ALE2 lineage contained representative convergent mutations, a clone from Flask 114 was chosen for further analysis. We will hereon refer to the ALE2-Flask114 strain as the evolved Nissle strain.

The evolved Nissle host displayed a differential improvement compared to the wildtype strain in the evolution growth media environment. We detected only a small improvement in growth in the evolved strain compared to the wildtype strain in the complex media without PQ. However, when increasing levels of PQ stress were introduced, a proportional reduction in growth for wildtype Nissle was observed whereas evolved Nissle was unchanged from the non-stress condition (Fig 2a, Fig S3b). As we noticed non-log-linear growth in batch culture which prevented accurate measurement of a single growth rate, likely due to multiple substrate utilization phases from the complex DMEM media, we chose to quantify the area-under-curve (AUC) of the growth curves. These findings show that the evolved strain did not exhibit a tradeoff in growth rate in the unstressed condition while conferring ROS stress resistance. Since the evolution conditions included both an alternate media and PQ-induced ROS stress, we chose to characterize their response to each condition individually.

**Fig. 2:**
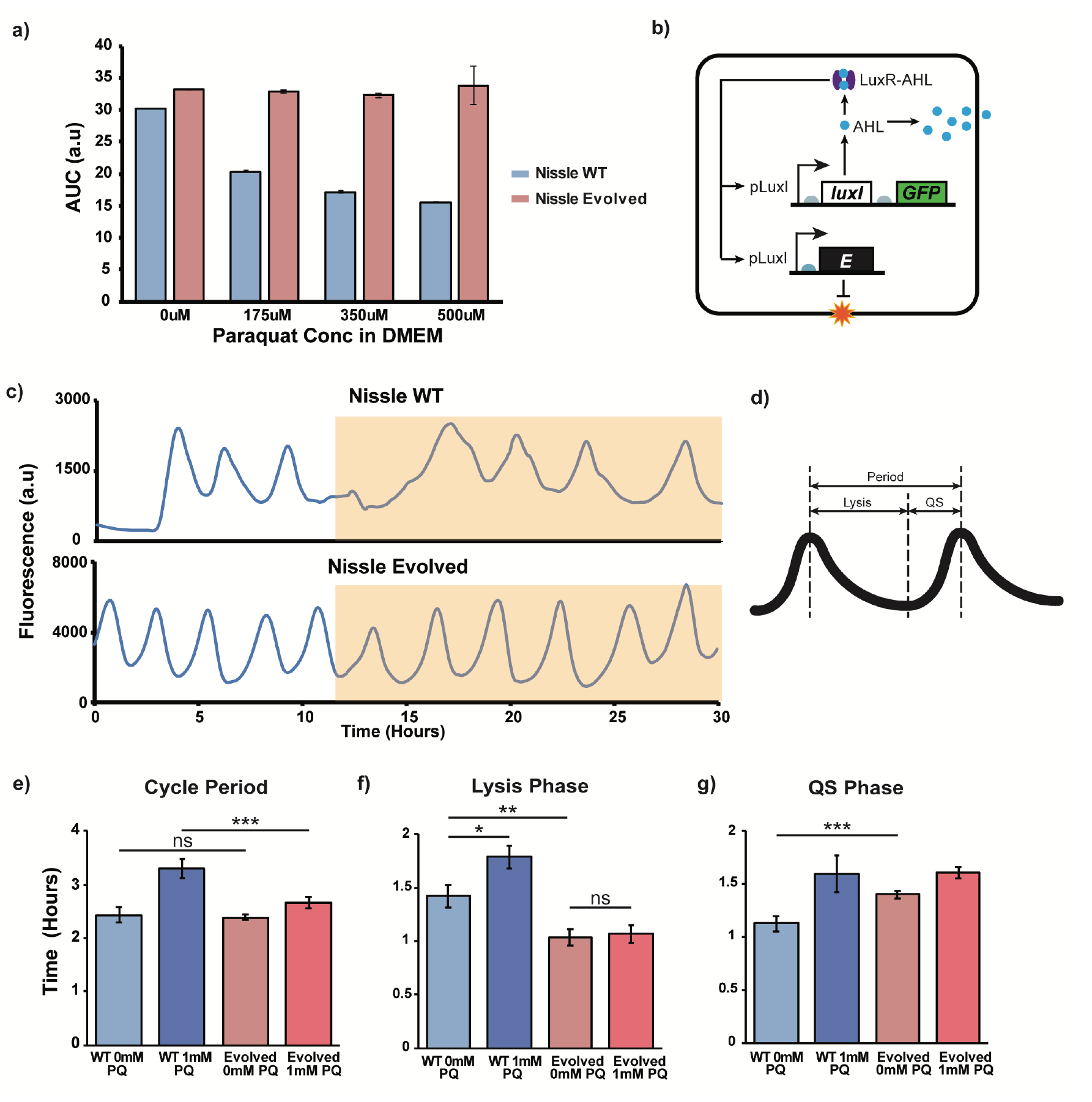
Characterization of wildtype and evolved Nissle strain response to ROS stress. **a)** Comparison of wildtype and evolved Nissle growth in DMEM-based evolution media in presence of increasing Paraquat concentrations. **b)** Diagram of two-plasmid SLC. In this version, luxI and GFP are on a high copy plasmid while X174E is on medium copy plasmid. **c)** Representative time traces of GFP fluorescence of wildtype and evolved Nissle with SLC growing in microfluidic traps in continuous culture of LB. Yellow shaded region indicates introduction of 1mM Paraquat. **d)** Schematic of SLC characterization. Each cycle of GFP oscillation is broken down into Lysis and Quorum Sensing (QS) phases for comparison. **e-g)** Comparison of cycle period, lysis phase period, and QS phase period for wildtype and evolved Nissle before and after 1mM Paraquat introduction in microfluidics. e) *P*_*ns*_ = 0.62 (*N*_*WT*_ = 14, *N*_*evolved*_ = 57), ^***^P = 0.0015 (*N*_*WT*_ = 16, *N*_*evolved*_ = 40). f) ^*^P = 0.0213 (*N*_0*mM*_ = 21, *N*_1*mM*_ = 19), ^**^P = 0.0062 (*N*_*WT*_ = 21, *N*_*evolved*_ = 60), *P*_*ns*_ = 0.7807 (*N*_0*mM*_ = 60, *N*_1*mM*_ = 44). g) ^***^P = 0.0008 (*N*_*WT*_ = 15, *N*_*evolved*_ = 60).

We first studied the effect of evolution to PQ on SLC dynamics in LB. We chose to characterize a two-plasmid version of the SLC that was previously used in *Salmonella* for delivery of cancer therapy as our candidate circuit (*3*). In this circuit, *pLux* driving X174E is on a medium copy p15A origin plasmid and *pLux* driving luxI and GFP are on a high copy ColE1 origin plasmid (Fig 2b). Previous work integrated the SLC into the Nissle genome for therapy delivery, however, its dynamics were not characterized in detail in microfluidics (*4*). We developed a two-plasmid version (SLC-EcN) that functions in Nissle (Table S2). After transforming into both wildtype and evolved Nissle, both strains were tested in the same multi-strain microfluidic device using LB with and without PQ (Fig 2c, Video S2). We saw similar oscillation cycle periods in both strains when growing in LB without PQ (Fig 2e). We took a closer look at the population dynamics by breaking each cycle into a lysis phase and quorum sensing (QS) phase for comparison. The lysis phase consists of the period from the highest GFP fluorescence to the lowest fluorescence due to population death, while the QS phase consists of the period from the lowest GFP fluorescence to the highest fluorescence due to AHL accumulation after lysis recovery (Fig 2d).

In LB without PQ, the lysis phase of the wildtype strain was significantly longer than that of the evolved strain, suggesting the wildtype strain was more affected by X174E and required longer recovery before restarting growth (Fig 2f). Testing in batch culture, where *pLux* only drives X174E, showed that after lysis induction with 10nM AHL, the wildtype strain exhibited less robust recovery compared to the evolved strain (Fig S4a, b). Thus, evolution to PQ-induced ROS is directly able to improve recovery of lysis survivors in Nissle. Meanwhile, in the QS phase, when grown without PQ, we noticed that the wildtype strain required significantly less time to reach quorum compared to the evolved strain (Fig 2g). In batch culture in LB, with *pLux* driving GFP, we saw similar GFP induction magnitudes between the two strains, suggesting both strains showed similar sensitivity to AHL (Fig S4c). Thus, we again attribute this difference to improved ROS tolerance in the evolved strain. Previous work shows that *E. coli*’s native aerobic response control system ArcAB, which is partially active, represses *pLux* (*38–40*). In the presence of oxidative stress, ArcAB is inactivated, relieving the repression. In the wildtype strain, there is less tolerance of ROS, suggesting inactivation of ArcAB, which increasingly activated *pLux* and reduced the amount of time needed to reach quorum. Meanwhile, mutations in the evolved strain increased ROS tolerance, leading to a more active ArcAB and less active *pLux*, resulting in a longer quorum sensing phase.

After 11 hours of growth, 1mM of PQ was added to the media. We chose to use 1mM PQ in order to place both strains under ROS stress, as the evolved strain seemed unaffected by 500uM PQ (Fig 2c). After PQ addition, the cycle period increased for both strains, but the wildtype strain oscillated at a significantly longer period compared to the evolved strain, suggesting that although both strains were affected by the ROS induced by PQ, the effect was reduced in the evolved strain (Fig 2e). We saw an increase in the QS phase for both strains but less so for the evolved strain (Fig 2g). Meanwhile, the lysis phase significantly increased in the wildtype strain whereas the evolved strain remained unaffected, further confirming that PQ evolution directly improved Nissle recovery from lysis component of the circuit (Fig 2f).

Our results demonstrate that evolution to PQ led to a significant improvement in circuit dynamics when exposed to ROS. Further, the mutations accumulated from this evolution directly influenced Nissle response to different SLC circuit components that produces or are influenced by ROS. In particular, we found the increased tolerance to X174E and subsequent reduction in lysis recovery period to be very promising, as it is an important component for therapy delivery in cancer applications.

### ALE In Combination with RBS Library Screening Improved Circuit Behavior in Complex Growth Media

Although we only observed a small improvement in Nissle growth rate in the complex DMEM-based media after evolution, we found a converged mutation related to metabolism of lysine and arginine (*argP*), both of which are found in the media (Table S1). We hypothesized that we would still see differences in SLC behavior between the wildtype and evolved strain in the complex media without PQ. However, in microfluidics, the evolved strain with SLC only exhibited one large lysis peak after which the cells were unable to recover, with only a few cells remaining in a fluorescent state (Fig 3e, Video S3). We attributed this to significant changes in growth media conditions, which may alter the metabolic needs of the cells and their response to different components of the circuit.

**Fig. 3:**
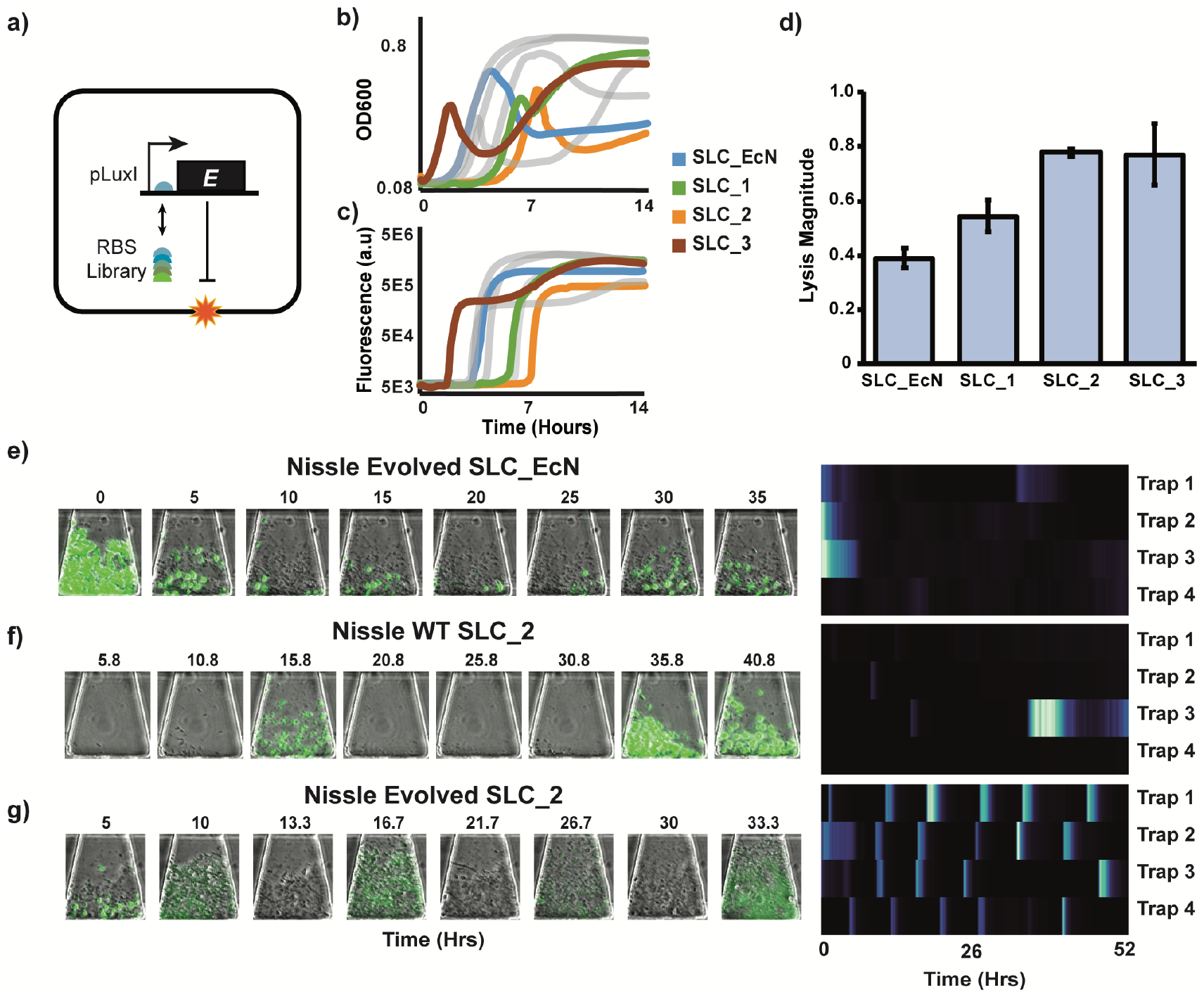
Optimization and characterization of wildtype and evolved Nissle circuit dynamics in complex media. **a)** Schematic of SLC library generation via directed mutagenesis of X174E protein ribosome binding site (RBS). **b-c)** Batch culture results of SLC library. Both OD600 and GFP fluorescence were characterized over 14 hours. Three SLC versions were chosen that represented the range of SLC behaviors. **d)** Comparison of lysis magnitudes associated with each RBS. Bars represent mean ± SEM (n = 3). **e)** Representative time series images from transmitted light and fluorescence channels of evolved Nissle with original SLC-EcN, wildtype Nissle with optimized SLC-2, and evolved Nissle with optimized SLC-2 in complex DMEM media. **f)** Corresponding heatmaps of GFP time series data for each strain.

We hypothesized that SLC dynamics in the complex media can be optimized by tuning the production of X174E (Fig 3a). Using a ribosome binding site (RBS) generator as well as sequences found in previous work, we generated a library of RBSs of X174E using directed mutagenesis (*3, 41*) (Table S2). In LB, growth in batch culture showed distinct dynamics in growth and GFP production (Fig 3b, c). Three SLC versions, SLC-1, SLC-2, and SLC-3 reflected different batch culture dynamics and were chosen for further analysis. To examine these RBSs and their potential effects on the SLC, we isolated the “lysis” part of the circuit by removing AHL production, in which each RBS is placed in front of X174E which is driven by *pLux*. Lysis magnitude was calculated for each variant at 10nM of AHL induction. SLC-EcN yielded the smallest lysis magnitude. SLC-1 showed the second strongest lysis, followed by SLC-2 and SLC-3 with similar lysis magnitudes (Fig 3d).

We found that when the evolved Nissle strain was transformed with SLC-2, we were able to again see consistent lysis dynamics (Fig 3f). The cycle period was significantly longer in the complex media, with a period of 8.93 +/- 0.72 hours compared to 2.38 +/- 0.05 hours in LB. Meanwhile, the wildtype strain with SLC-2 exhibited only 1 or 2 lysis peaks after which it was unable to recover in growth. Further, the wildtype strain had trouble growing in the microfluidic traps, most likely due to burden imposed by the circuit (Fig 3g). We confirmed this in batch culture in the complex media, again using *pLux* driving X174E. After 10nM AHL induction of lysis, growth recovery was less robust in the wildtype strain after normalizing by growth without lysis (Fig S5b). Further, the wildtype strain harboring SLC-2 took a significantly longer time to begin growth compared to the evolved strain (Fig S5c). Finally, when we studied the response of each strain to AHL using *pLux* driving GFP, we saw that at 10nM AHL induction, the evolved strain produced more GFP compared to the wildtype (Fig S5d).

Thus, although we observed only a small difference in growth rate between the wildtype and evolved strains in batch culture, the accumulated mutations from ALE improved growth and behavior with the SLC. Further, our results show that although ALE ensures improved adaptation to complex growth environments, circuit parameters may also need to be re-tuned for robust circuit dynamics.

### Comparison of Nissle and MG1655 Circuit Behavior

During our investigation, we noticed different circuit responses between Nissle and MG1655 due to inherent genomic differences between the two strains. Previous comparative studies between Nissle and other *E. coli* strains showed that Nissle shared only 73.2% of homologous genes with MG1655 (*42*). Although the growth behavior of Nissle has been well-studied through metabolic network reconstruction, little work has been done to compare synthetic circuit response between Nissle and MG1655 (*43*). Therefore, we sought to gain further insight in the differences in circuit behavior between wildtype MG1655 and Nissle by focusing on SLC dynamics. Since MG1655 grew poorly in the DMEM-based media, we performed our comparisons in LB.

Wildtype Nissle strains harboring SLC variants developed for screening in DMEM-based media were examined in further detail in a microfluidic chip (Table S2). Compared to consistent oscillations seen in SLC-EcN, an increase in RBS strength in SLC-1 led to no growth or growth without oscillations. Further increase in RBS strength (SLC-2) led to more variability between traps, with one or two lysis events before loss of oscillations. SLC-3 yielded similar dynamics to SLC-2, with the population generally exhibiting one strong lysis event followed by weak or complete loss of oscillatory behavior after regrowth (Fig 4a). We next transformed the same SLC variants into wildtype MG1655. As expected, MG1655 exhibited significantly different behavior to each SLC variant compared to Nissle. Whereas SLC-EcN yielded consistent oscillations in Nissle, we saw extremely weak or no oscillations in MG1655 followed by continuous growth in the traps. Meanwhile, increase in RBS strength in SLC-1 led to consistent oscillations in all traps. All traps of SLC-2 showed one strong lysis event with two traps showing two lysis events before loss of oscillations. In SLC-3, one strong lysis event followed by loss of oscillatory behavior was observed. Overall, we observed that MG1655 was able to oscillate at stronger RBS strengths (Fig 4a).

**Figure 4.**
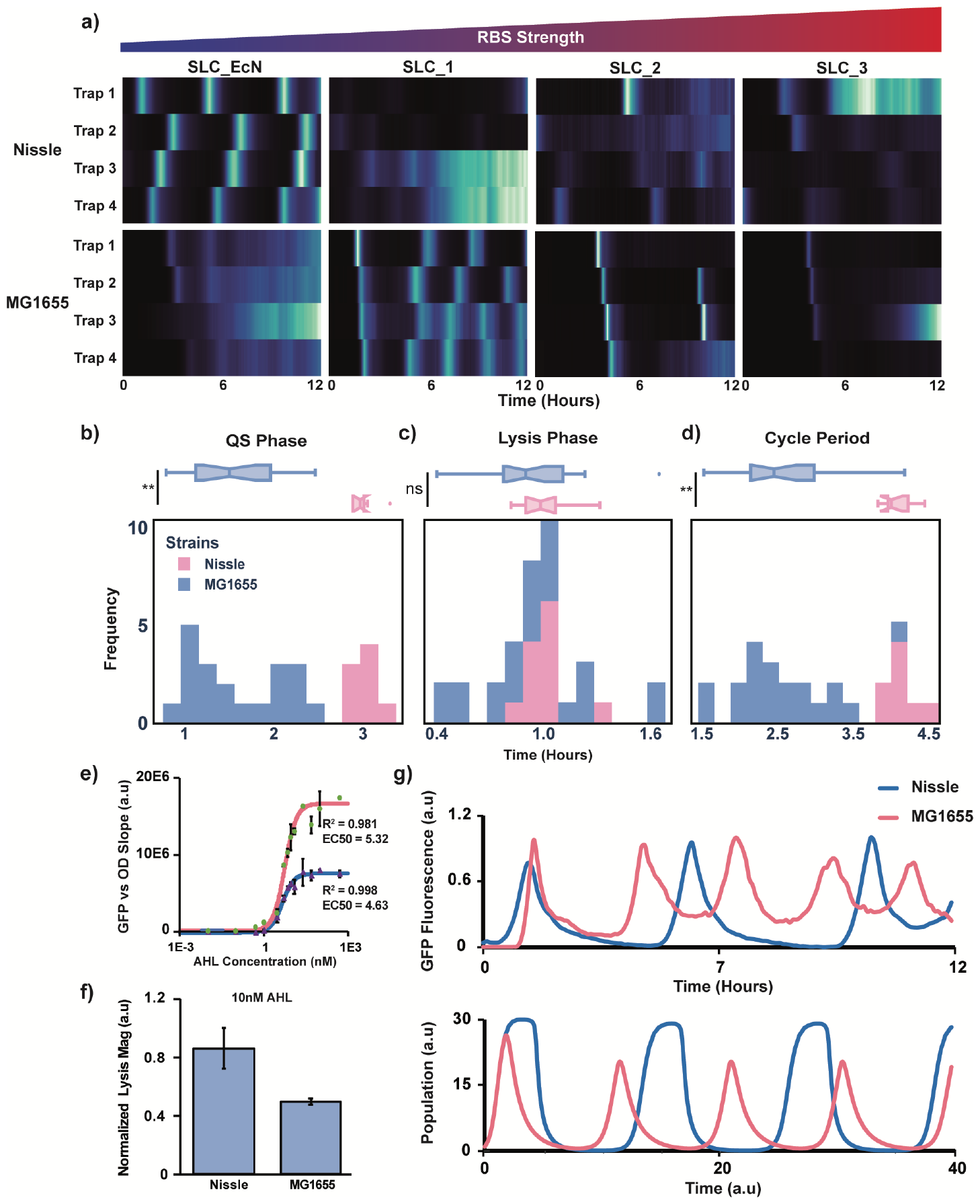
Characterization of wildtype Nissle and MG1655 SLC population dynamics. **a)** Heatmaps of GFP time series data of wildtype Nissle and MG1655 with different SLC RBS library versions in microfluidics. **b-d)** Comparison of QS phase, lysis phase, and cycle period of Nissle and MG1655. Top: Boxplots of each period distribution. Bottom: Histogram of each period distribution. b) ^**^P = 0.0001 (*N*_*Nissle*_ = 8, *N*_*M*_ *G*1655 = 20). c) *P*_*n*_*s* = 0.5319 (*N*_*N*_ *issle* = 12, *N*_*M*_ *G*1655 = 24). d) ^**^P = 0.0001 (*N*_*N*_ *issle* = 8, *N*_*M*_ *G*1655 = 20). **e)** Dose-response curve of AHL vs GFP induction response. Data points represent mean ± SEM (n = 3). Solid line represents fitted Hill function. **f)** Comparison of lysis susceptibility between MG1655 and Nissle. Bars = mean ± SEM (n = 3). **g)** Top: representative time series GFP data of MG1655 and Nissle SLC dynamics. Bottom: Computer simulations of mathematical model for each strain.

We next compared the dynamics between Nissle with SLC-EcN and MG1655 with SLC-1 by quantifying each part of the SLC cycle. We found that the QS phase in MG1655 was significantly shorter compared to Nissle, contributing to a significantly faster lysis cycle (Fig 4b, d). We examined the responsiveness of the *pLux* promoter of each strain by using a gradient of AHL to induce *pLux* driving GFP expression. We quantified the slope of the OD vs GFP plots at log growth following induction to get an estimate of GFP production per cell. After fitting to a Hill function, we saw that both strains had similar EC50s, suggesting that *pLux* in both strains have similar sensitivity. However, we saw that GFP expression level of MG1655 was consistently higher (two-fold higher response at 10nM AHL induction), suggesting that for the SLC, MG1655 has a higher luxI production rate than Nissle, leading to a faster accumulation of AHL and a shorter time to reach quorum threshold (Fig 4b, e). When we compared the Lysis Phase, we saw no significant difference between the two strains (Fig 4c). We next examined each strain’s susceptibility to the X174E protein using *pLux* promoter driving X174E instead of GFP. After induction with 10nM AHL, we saw the lysis magnitude of Nissle was higher than that of MG1655 after normalizing by protein production (Fig 4f). Using a simple deterministic model of SLC dynamics adapted from previous works (*3,44*), we were able to replicate the SLC dynamics that were observed in each strain (Fig 4g). We confirmed that period and QS Phase correlates negatively with increased lysis sensitivity while the Lysis Phase positively correlates.

Meanwhile, all three metrics correlates negatively with increased AHL production rate (Fig S6). Thus, similar Lysis Phases between the MG1655 and Nissle may be due to a balance between increased AHL and reduced lysis sensitivity in MG1655. Our results demonstrate that inherent genomic differences play an important role in host-circuit response.

## Discussion

Despite continuous development of tools for rational engineering of synthetic circuits (*45–47*), the need for trial-and-error testing and mass screening remains a challenge in the creation of complex circuits (*8, 14*). This problem is further exacerbated in the move towards real-world deployment, where circuits developed in the lab must now function predictably in poorly characterized and new growth environments. This study demonstrated the feasibility of combining ALE and traditional circuit engineering approaches for the transition from lab-originated design to translational application, where an intricate circuit can be made to regain function in a complex growth environment. In M9 minimal media with lactate, MG1655 was able to regain SLC dynamics through host evolution alone. Meanwhile, evolution in a complex DMEM-based media not only significantly reduced the effect of Paraquat-induced ROS stress on Nissle, but also improved its tolerance of burdensome SLC components. In combination with directed mutagenesis of the SLC, the evolved Nissle with SLC was able to fully oscillate in the complex media environment. While a number of previous studies have mainly focused on using adaptive laboratory evolution for improving growth rates of bacteria and yeast for bioproduction, this work leverages ALE’s power for robust synthetic circuit function for potential therapeutic applications. Further, as both *E. coli* Nissle and quorum sensing-based systems are becoming increasingly popular in clinical applications, this work provides a detailed comparison and modelling of SLC dynamics between MG1655 and Nissle.

The methods presented in this study can be applied to the development of a wide variety of circuits in a range of host strains given the generalizability of evolutionary engineering techniques. One immediate example is in the realm of bacterial cancer therapy, where strains such as *E. coli*, and *Salmonella typhimurium* can potentially be anaerobically evolved for improved growth, circuit response, and therapy delivery in the hypoxic tumor core (*48, 49*). While the genotypes of the evolved optimized clones were determined, further work is necessary to understand the impact of the specific mutations on the overall phenotype. Such studies are informative, but often require significant reverse engineering and omics analyses to elucidate mutational mechanisms (*50*). Overall, this study provides a framework for the incorporation of evolutionary engineering and rational synthetic circuit design for the development and eventual deployment of complex genetic circuits.

## Acknowledgments

J.T.Z., A.L., P.E., and J.H. are supported by the National Institute of Biomedical Imaging and Bioengineering Grant No. R01 EB030134. J.T.Z. and A.L. are also supported by the National Science Foundation Graduate Research Fellowship under Grant No. DGE-2038238. Any opinions, findings, and conclusions or recommendations expressed in this material are those of the author(s) and do not necessarily reflect the views of the National Science Foundation. C.A.O., A.M.F., and M.W were funded by the Novo Nordisk Foundation grant NNF20CC0035580 and by the National Institute of General Medical Sciences grant GM057089. The authors thank Nicholas Csicsery for providing the multi-strain microfluidic device, Aayush Somani for assistance with batch culture experiments, and Robert Cooper for critical reading of the manuscript.

## Supplementary materials

Materials and Methods

Figs. S1 to S6

Tables S1 to S3

References (*14, 41, 51*)

Movies S1 to S3

