## Supplementary material for "Host evolution improves genetic circuit function in complex growth environments": Materials and Methods, Supplemental Figs 1-6, Supplemental Table 2-3

### Supplementary Information

### Materials and Methods

**Adaptive Laboratory Evolution** ALE was performed in 15mLs of evolution media at 37C and well-mixed at 1100 rpm for full aeration. An automated system passed the culture to fresh flasks before they had reached stationary phase (Tecan Sunrise plate reader, equivalent to OD600 = 1 on a traditional spectrophotometer with 1cm path length). A maximum specific growth rate ($h^{-1}$) was calculated by linear regression of ln(OD600) and time (h) during the exponential growth phase (OD600 = 1). For *E. coli* MG1655, the evolution media consisted of M9 minimal media with 2g/L lactate (Sigma, CAT 60389). Six replicates were evolved in parallel for identification of common mutation targets. For *E. coli* Nissle, the evolution media consisted of DMEM (Sigma, CAT D5030), 30mM lactate (Sigma, CAT 60389), 0.4mM glucose, 70mM MES (Sigma, CAT 69889), 1g/L NH4Cl, 350uM Paraquat (Thermo, CAT AC227320050), with pH adjusted to 6.4 using NaOH or HCl as needed. Five replicates were evolved in parallel.

**DNA sequencing and mutation calling** DNA from each replicate at the endpoint of the evolution lineage was isolated from an overnight culture and centrifuged for 5 min at 8000rpm. The cell pellet was frozen at 80C. Genomic DNA was isolated using the Quick-DNA Fungal/Bacterial Microprep Kit (Zymo Research) following manufacturer's protocol, including RNAse A treatment. Sequencing libraries were prepared using a Kapa Hyper Plus Kit (Roche Diagnostics) following manufacturer's protocol. Libraries were run on HiSeq and/or NextSeq (Illumina). Sequencing reads were filtered and trimmed using AfterQC version 0.9.7.75. We mapped reads to the *E. coli* K-12 MG1655 reference genome (NC_00913.3) and *E. coli* Nissle reference genome (NZ_CP022686) and cryptic plasmids (NZ_CP058219, NZ_MW240712) primarily using breseq version 0.33.1.5 as a part of ALEdb's mutation calling platform (*52*). Mutation analysis was performed using ALEdb website 55 (*51*).

**Plasmids and Strains** MG1655 and Nissle strains (wildtype and evolved) were cultured either in lysogeny broth (LB) media or evolution media. Appropriate antibiotics were added for plasmid maintenance. Plasmids pSpSLC_RBS11, pTD103-sfGFP, and pZA-X174E-pTacHlyE were constructed by our group in previous studies. pZA-X174E-EcN was obtained by replacing the ribosome binding site of X174E of pZA-X174E-pTacHlyE (Table S2). Plasmids were transformed into all MG1655 strains via chemical transformation, and Nissle strains via electroporation (Table S3).

**Library creation for Evolved Nissle SLC strains** *E. coli* Nissle A2-F114 was first made electrocompetent, transformed with pTD103-sfGFP via electroporation, and plated on LB agar plate containing kanamycin. The next day, one colony of Nissle containing pTD103-sfGFP was selected, grown up overnight, and made electrocompetent.

To generate a mutant library of the pZA-X174E-EcN plasmid, a ribosome binding site library calculator was used for site-directed mutagenesis primer design (*41*). The entire plasmid was PCR amplified with the following degenerate primers where N indicates any base: 5' TAATAGATATAAATANGGANNTTGNC ATGGTACGCTGGACTTTG 3' and 5' ATTCGACTATAACAAACCATTTTC 3'. The PCR reaction was incubated with DPNI at 37C for 1 hour to digest template plasmid. The resulting PCR product (3.1kb) was run on an agarose gel and extracted using a QIAquick Gel Extraction Kit (QIAGEN). Blunt-end DNA ligation was performed by mixing 4uL of the gel-extracted PCR product with 0.5uL T4 ligase buffer, 0.5uL T4 PNK, 1uL of DNase-free water and incubated at 37C for one hour. Next, 0.5uL T4 ligase buffer, 0.5uL T4 DNA ligase, and 4uL of DNase-free water were added to the reaction mixture and incubated at room temperature overnight. The next day, 50uL of chemically competent *E. coli* DH5alpha cells were transformed with 5uL of the reaction mix and plated on an LB agar plate containing 0.2% glucose and chloramphenicol. The entire plate was scraped into 5mL LB to harvest all colonies, and pooled library plasmids were isolated using the QIAprep Spin Miniprep Kit. Following sequencing confirmation of library creation, the pooled library plasmids were electroporated into electrocompetent *E. coli* Nissle strains containing pTD103-sfGFP and plated on an LB agar plate containing 1% glucose, kanamycin, and chloramphenicol for 15 hours. 48 colonies from the agar plate were randomly selected for mutant screening and grown up for 16 hours in LB media with 1% glucose, kanamycin, and chloramphenicol prior to use in experiments. 2uL of each strain was added to 200uL of fresh media with antibiotics in a standard Falcon tissue culture 96-well flat bottom plate. Cells were incubated at 37C shaking in a a Tecan Infinite M200 Pro. The OD at 600nm absorbance and GFP fluorescence (488nm excitation, emission 520nm) was measured every 5 mins.

**Multi-strain microfluidic device fabrication and experimental protocol** A poly-dimethylsiloxane (PDMS) device was made from the microfluidic device wafer by mixing 77 grams of Sylgard 184 and pouring it on the wafer centered on a level 5”x5” glass plate surrounded with an aluminum foil seal. The degassed wafer and PDMS was cured on a flat surface for one hour at 95C. We used a multi-strain microfluidic device that was previously developed in the lab (*14*). The device consists of a 6 x 8 array of cell-trapping regions that can be loaded with liquid bacterial cultures. Each position contains four smaller cell traps downstream of the large trapping region that serve as regions of interest (ROIs) for tracking population dynamics in fluorescent and transmitted light channels. Up to 48 distinct positions can be loaded with a unique *E. coli* strain for continuous culture and data acquisition at high spatiotemporal resolutions.

Cells were grown overnight on LB media with appropriate antibiotics and 1% glucose to suppress expression of the *pLux* promoter driving lysis. 200uL of each strain was spun down and resuspended in 50uL of media with 0.075% Tween-20. A PDMS device and 4” x 3” glass slide were cleaned with 70% Ethanol and exposed to oxygen plasma. Each strain was transferred to a cell-trapping region via handspotting using a 30 gauge syringe tip. The device and glass slide were bonded together and cured at 37C for two hours. Following curing, the device was placed in a vacuum for 30 minutes, then mounted onto a custom optical enclosure. The inlet port was connected to a 30mL syringe and tygon tubing with 15mL of LB media, antibiotics, and 0.075% Tween-20 (used as surfactant to reduce clogging). The waste port was connected to a 30mL syringe and tygon tubing with 1mL of MilliQ water. The height difference between inlet and outlet was 2” corresponding to a flow rate of approximately 0.5mLs/hr. After traps were filled to confluence, LB media was removed in entirety and replaced with 15mL of evolution media containing appropriate antibiotics and 0.075% Tween-20.

**Imaging and data extraction** Images were acquired with a Nikon TI2 using a Photometrics CoolSnap cooled charge-coupled device (CCD) camera. The microscope was housed in a plexiglass incubation chamber maintained at 37C by a heating unit. Phase-contrast images were taken at 10X magnification at 100 $\mu$s exposure time. Fluorescent exposure times were 200 $\mu$s at 30% intensity for GFP. Images were taken every 5 mins for every experiment. Fluorescence intensity profiles were extracted by analyzing frames from the GFP fluorescent channel. Images were processed in ImageJ, where a custom ROI manager was used to extract mean fluorescence values within each trap. Fluorescence values were normalized by subtracting the local background signal.

**SLC cycle analysis** After data extraction, functions within the scipy signal processing toolbox were used. Fluorescence traces were first smoothed using the savgol_filter. Peaks and troughs were then identified using the argrelextrema function in the scipy signal processing toolbox, filtering for peaks and troughs with a minimum prominence of 8 (arbitrary fluorescence units). Within each cycle, cycle period was calculated as the peak-to-peak time, growth phase was calculated as previous trough subtracted by current peak time, lysis phase was calculated as current peak subtracted by following trough time.

**SLC library transformation into wildtype Nissle and MG1655 strains** SLCs of interest were identified in the evolved Nissle strains via batch culture and microfluidic analysis. The entire pZA-X174E-Lib plasmid in each SLC strain of interest were PCR amplified using the following primers: 5' AATCGCCAGCGGCATCA 3' and 5' CAGGTTCATCATGCCGTCTGT 3'. The PCR reaction was processed and re-ligated as described in the "Library creation for Evolved Nissle SLC strains" section. The re-ligated PCR was transformed into DH5alpha and plated on LB agar with chloramphenicol and 0.2% glucose. The next day, one colony was selected and grown up overnight for plasmid extraction. Following sequencing confirmation, the pZA-X174E-Lib plasmid was co-transformed with pTD103-sfGFP into wildtype Nissle or MG1655.

**Batch culture experiments** For comparison of wildtype and evolved strain response to evolution media, the appropriate strains were seeded from a -80 C glycerol stock into 3mL of the evolution media and appropriate antibiotics, and incubated in a 37C shaking incubator. The following day, 2uL of overnight culture were added to 200uL of fresh evolution media with appropriate antibiotics and Paraquat in a standard Falcon tissue culture 96-well flat bottom plate. Cells were incubated at 37C shaking in a Tecan Infinite M200 Pro. The OD at 600nm absorbance and GFP fluorescence (488nm excitation, emission 520nm) was measured every 5 mins. For Nissle growth rate calculation, the area-under-curve (AUC) of the OD600 curve was calculated for the first 10 hours of growth.

**GFP induction calculation** A plasmid with *pLux* promoter driving GFP expression (M4) was transformed into strains of interest. 2uL of each strain was added to 200uL of fresh evolution media or LB with spectinomycin and desired AHL concentration. Each well was monitored in a Tecan plate reader for 12 hours for both OD600 and GFP. OD600 vs GFP was plotted for each well. For calculating GFP expression of MG1655 in M9 lactate (Fig S2), AUC of OD vs GFP curves was measured. The ratio of AUCs between 1nM and 0nM AHL induction was calculated for each strain to get the relative AUC for comparison. For calculating GFP expression of Nissle, the OD vs GFP between OD600 of 0.4 – 0.55 was fitted to a linear regression and the slope was calculated. For characterizing pLux dose-response between MG1655 and Nissle (Fig S7f), AHL concentrations between 0.01nM to 500uM was added and growth response was monitored as described. A dose response curve was generated with the OD-GFP slope vs. AHL concentration by fitting the data to the Hill Equation.

**Lysis magnitude calculation** To experimentally determine relative expression strength of RBS sequences from the SLC library (Fig 3d), pZA-X174E-Lib plasmids were isolated from each SLC strain of interest as described in "SLC library transformation into wildtype Nissle and MG1655 strains" section. These plasmids consist of pLux promoter driving X174E, where lysis is inducible by AHL. After confirmation by sequencing, each plasmid was transformed into E. coli DH5alpha and plated on LB agar with 0.2% glucose and chloramphenicol. One clone was isolated and grown up overnight. 2uL of each overnight culture was added to 200uL of fresh LB with chloramphenicol in triplicates in a 96-well flat bottom plate and monitored for OD600 in a Tecan microplate reader at 37C with orbital shaking. Once cultures reached OD 0.3, the well plate was quickly removed from the microplate reader and AHL was added to achieve a final concentration of 10nM. The well plate was then re-inserted into the Tecan and cultures were grown for 12 hours. The growth curve for each well was examined for the first inflection point where the derivative of the culture OD with respect to time changed from positive to negative. The lysis magnitude was calculated as L/G, where G is the positive change in OD from the initial time point to the inflection point and L is the negative change in OD from the inflection point to the time point at which OD was lowest following the inflection point. To compare susceptibility to the X174E between MG1655 and Nissle (Fig S7f), the lysis magnitude of Nissle at 10nM AHL was normalized by multiplying with $GFPInduction_{MG1655}/GFPInduction_{Nissle}$ where GFP induction is calculated as in "GFP induction calculation" section at 10nM AHL induction.

**Post-lysis growth calculation** Wildtype and evolved Nissle strains were transformed with pZA-X174E-pTacHlyE. Lysis was induced and monitored as described in "Lysis magnitude calculation". The growth curve for each well was examined for the lowest OD point following the first inflection point. The AUC from the lowest OD point to six hours after that point was calculated for post-lysis growth.

**Modeling** We constructed a simple deterministic model of the SLC that was adapted from previously published models of SLC dynamics. This model consists of three ordinary differential equations (ODEs) that describes cell growth and lysis-induced cell death (Equation 1), AHL production and accumulation (Equation 2), and X174E Lysis gene production (Equation 3). Equation 4 ($P_{lux}$) describes dynamics of the pLux promoter, which is brought to the ON state when AHL reaches a threshold. Equation 5 ($\gamma_N$) describes dynamics of cell-induced death from the lysis gene. Due to cellular growth and lysis occurring at a much slower time scale than delays in gene expression, we did not include a delay term in the model that accounts for delays in transcription and translation of LuxI and lysis gene. The parameters used in the model are: $\mu_G$ = 0.1 (dilution due to cell growth), $N_0$ = 30 (cell capacity of the trap), $b$ = 0.4 (basal AHL production), $\gamma_A$ = 2 (AHL degradation and dilution rate), $c_L$ = 3 (lysis gene copy number), $\gamma_L$ = 0.5 (lysis gene degradation and dilution rate), $m$ = 2 (Hill coefficient of pLux promoter), $kA$ = 5 (AHL binding affinity), $n$ = 2 (Hill coefficient of lysis protein), $k$ = 4 (maximum rate of cell lysis).

$$\frac{dN}{dt} = \mu_G N(N_0 - N) - \gamma_N N \tag{1}$$

$$\frac{dA}{dt} = b + \alpha * NP_{lux} - \gamma_A A \tag{2}$$

$$\frac{dL}{dt} = c_L P_{lux} - \gamma_L L \tag{3}$$

$$P_{lux} = \frac{A^m}{A^m + kA^m} \tag{4}$$

$$\gamma_N = k * \frac{L^n}{L_0^n + L^n} \tag{5}$$

To account for differences in AHL production rate, we varied $\alpha$ (AHL production rate) between 2.5 to 12.5. To account for differences in lysis gene susceptibility, we varied $L_0$ (concentration of lysis gene resulting in half-maximum lysis) between 0.02 to 2. To generate our final curves for MG1655, we chose $\alpha$ = 12 and $L_0$ = 2. For Nissle, we chose $\alpha$ = 3.11 and $L_0$ = 0.34. All model results were obtained in MATLAB using the ode23 function.

### Supplementary Figures

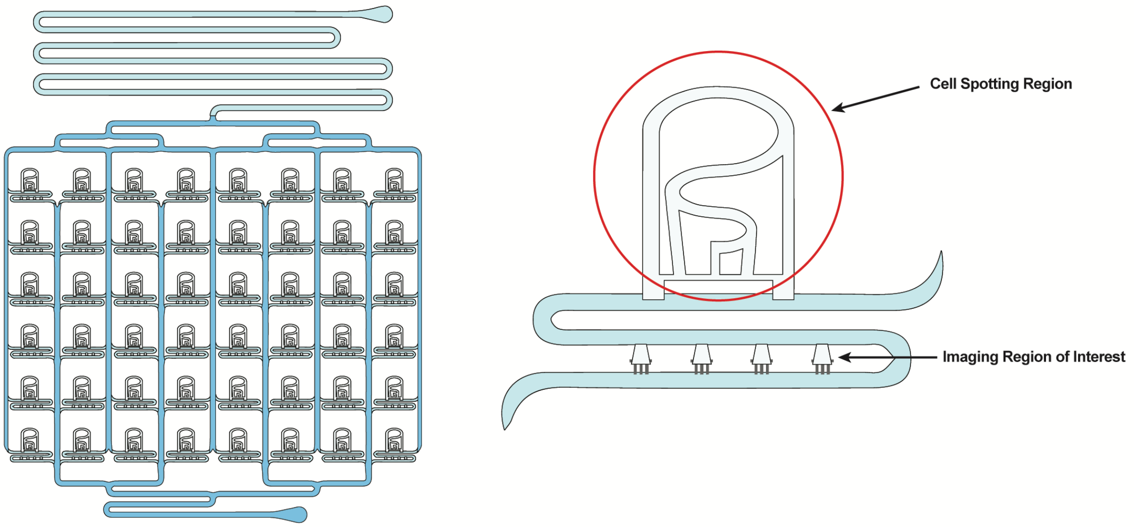


**SI Figure 1.** Device layout of a previously developed multi-strain microfluidic chip (*14*) The device consists of 6 by 8 array of cell-trapping regions that can be loaded with liquid bacterial cultures via hand-spotting. Four smaller cell traps downstream serve as regions of interest for tracking.

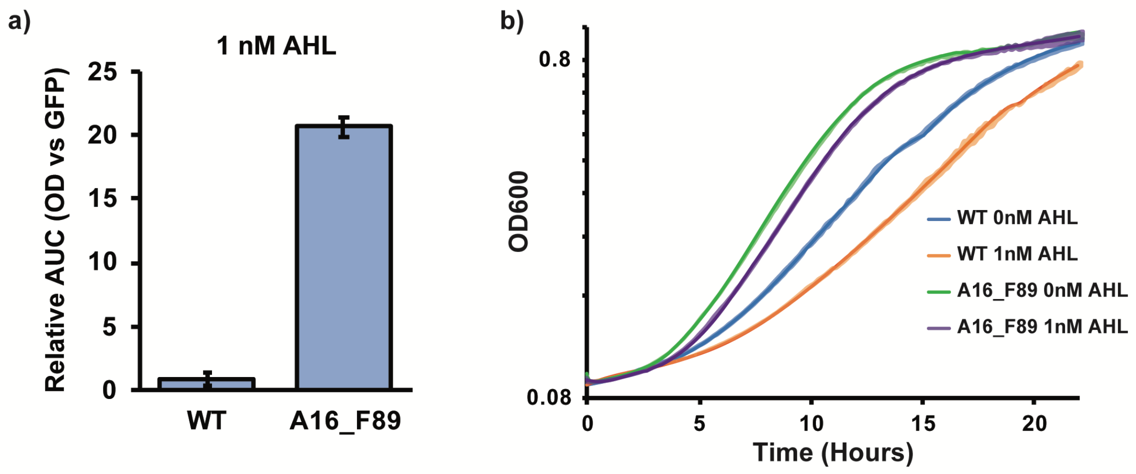


**SI Figure 2.** Characterization of MG1655 wildtype and A16-F89 strain response to GFP induction using 10nM AHL. **a)** Comparison of GFP induction at 1nM AHL addition. Bars represent mean ± SEM (n = 3). **b)** Comparison of OD600 growth curves with 1nM AHL addition. All data points represent mean (solid line) ± SEM (shaded areas), n = 3.

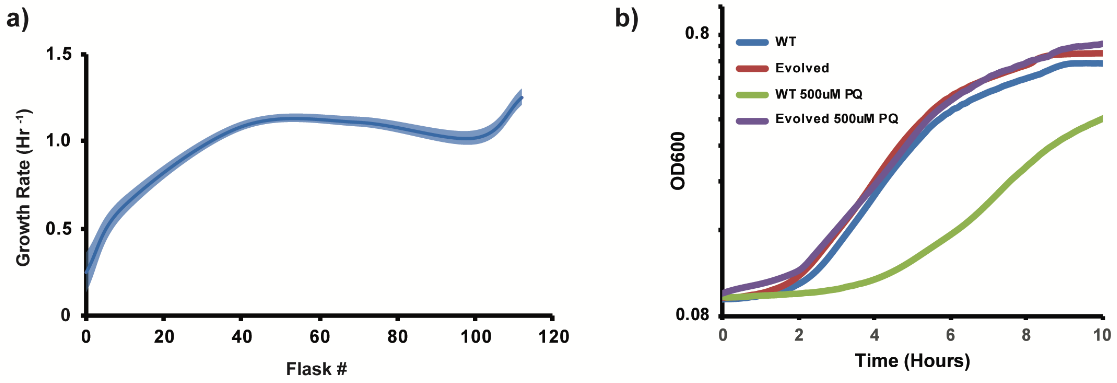


**SI Figure 3.** Characterization of Nissle growth during ALE. **a)** Growth rate of Nissle over course of ALE in complex media. All data points represent mean (solid line) ± SEM (shaded areas), n = 5. **b)** Growth curve comparison of wildtype and evolved Nissle in LB with and without 500uM Paraquat. All data points represent mean (solid line) ± SEM (shaded areas), n = 3.

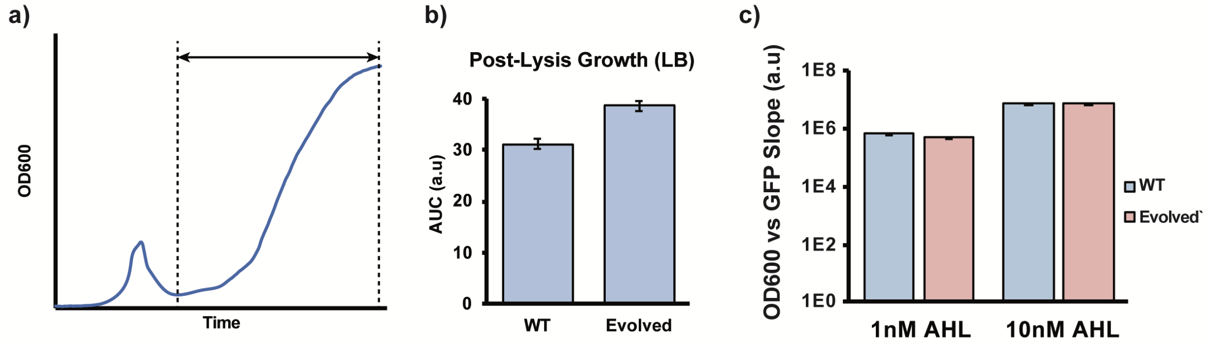


**SI Figure 4.** Batch culture characterization of lysis recovery and quorum sensing in wildtype and evolved Nissle in LB. **a)** Diagram of lysis recovery quantification. The AUC from the lowest OD point to six hours after that point was calculated for post-lysis growth. **b)** Comparison of post-lysis growth in LB. c) Comparison of GFP production after AHL induction. Bars represent mean ± SEM (n = 3).

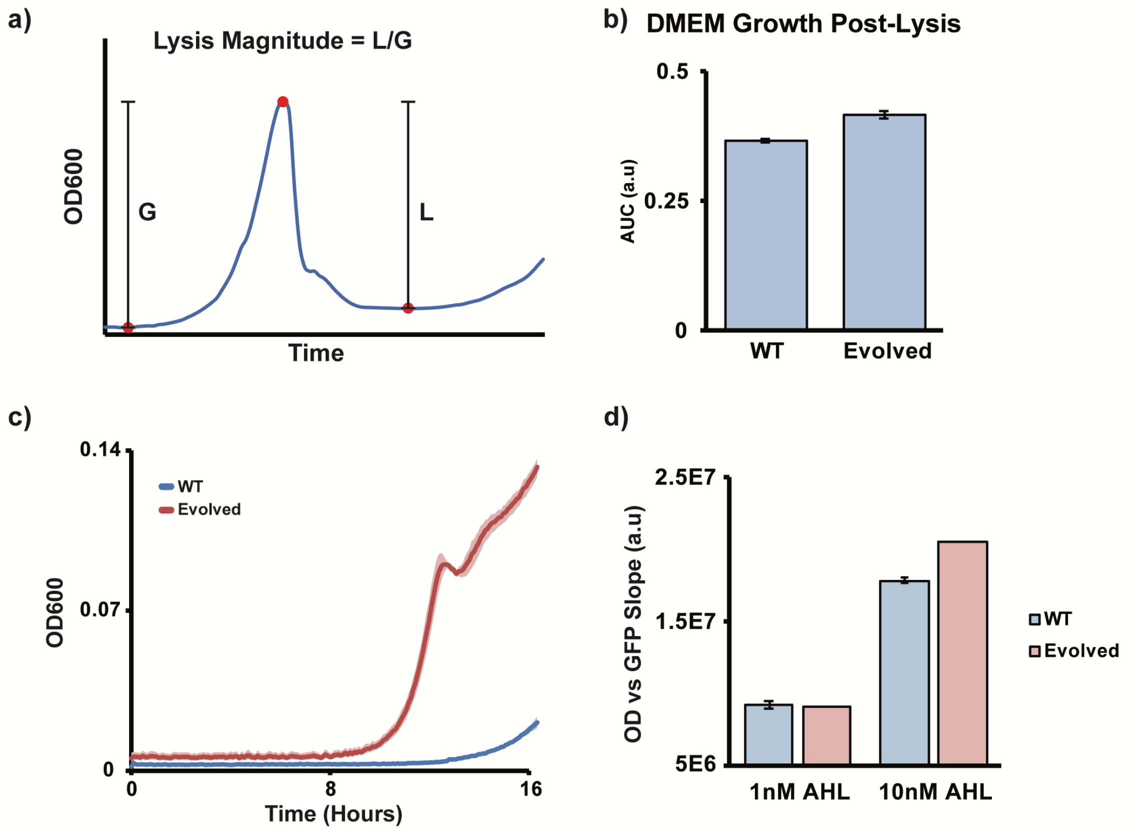


**SI Figure 5.** Batch culture characterization of wildtype and evolved Nissle in DMEM-based evolution media. **a)** Diagram of lysis magnitude calculation for different RBS strengths. **b)** Comparison of post-lysis growth. Bars represent mean ± SEM (n = 3). **c)** Comparison of batch culture growth with SLC-2. All data points represent mean (solid line) ± SEM (shaded areas), n = 3. **d)** Comparison of GFP production after AHL induction. Bars represent mean ± SEM (n = 3).

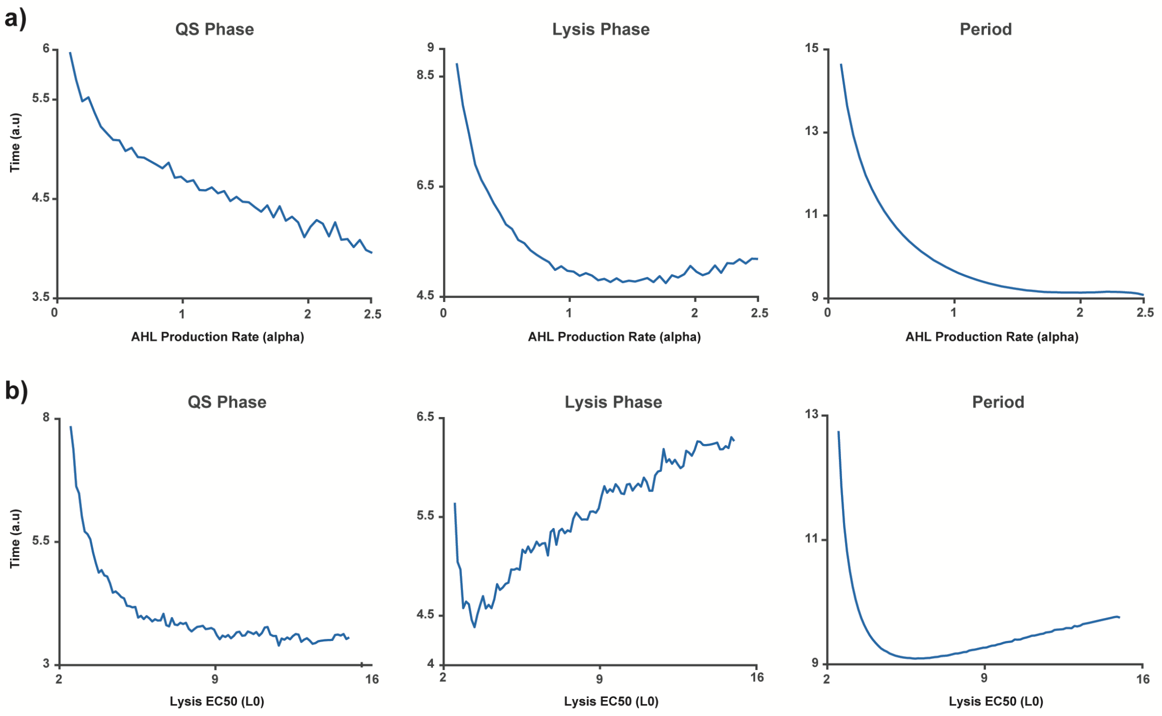


**SI Figure 6.** Simulation of lysis dynamics. **a)** QS, lysis, and cycle period changes with respect to increasing AHL production rate. **b)** QS, lysis, and cycle period changes with respect to lysis susceptibility (EC50).

Supplementary Table 2 – X174E RBS sequences for each 2-plasmid SLC version used in this paper

| SLC Version | X174E RBS Sequence |
|---|---|
| SLC_EcN | TAATAGATATAAATAAGGACCTTGAC |
| SLC_1 | ATAGATATAAATAAGGACATTGTC |
| SLC_2 | TAATAGATATAAATAAGGAATTTGAC |
| SLC_3 | AAACGCAAGGGAGGTTGGT |

Supplementary Table 3 - Strains and Plasmids

| Host Strain | Plasmid(s) | Figures |
|---|---|---|
| *E. coli* MG1655 WT | | 1 |
| *E. coli* MG1655 A16_F26 | | 1 |
| *E. coli* MG1655 A16_F51 | | 1 |
| *E. coli* MG1655 A16_F89 | | 1 |
| *E. coli* Nissle WT | | 2, SI3 |
| *E. coli* Nissle A2_F114 | | 2, SI3 |
| *E. coli* MG1655 WT | pSpSLC_RBS11 | 1 |
| *E. coli* MG1655 A16_F26 | pSpSLC_RBS11 | 1 |
| *E. coli* MG1655 A16_F51 | pSpSLC_RBS11 | 1 |
| *E. coli* MG1655 A16_F89 | pSpSLC_RBS11 | 1 |
| *E. coli* MG1655 WT | M4 | SI2 |
| *E. coli* MG1655 A16_F89 | M4 | SI2 |
| *E. coli* MG1655 WT | pTD103-sfGFP + pZA-X174E-RBS1 | 4 |
| *E. coli* Nissle WT | pTD103-sfGFP + pZA-X174E-EcN | 2, 4 |
| *E. coli* Nissle A2_F114 | pTD103-sfGFP + pZA-X174E-EcN | 2, 3 |
| *E. coli* Nissle WT | M4 | SI4, SI5 |
| *E. coli* Nissle A2_F114 | M4 | SI4, SI5 |
| *E. coli* Nissle WT | pZA-X174E-pTacHlyE | SI4, SI5 |
| *E. coli* Nissle A2_F114 | pZA-X174E-pTacHlyE | SI4, SI5 |
| *E. coli* Nissle WT | pTD103-sfGFP + pZA-X174E-RBS2 | 3, SI5 |
| *E. coli* Nissle A2_F114 | pTD103-sfGFP + pZA-X174E-RBS2 | 3, SI5 |
| *E. coli* DH5alpha | pZA-X174E-EcN | 3 |
| *E. coli* DH5alpha | pZA-X174E-RBS1 | 3 |
| *E. coli* DH5alpha | pZA-X174E-RBS2 | 3 |
| *E. coli* DH5alpha | pZA-X174E-RBS3 | 3 |
